# Presence of WH2 like domain in VgrG-1 toxin of *Vibrio cholerae* reveals the molecular mechanism of actin cross-linking

**DOI:** 10.1101/226605

**Authors:** Priyanka Dutta, A.S. Jijumon, Mohit Mazumder, Drisya Dileep, Asish K. Mukhopadhyay, Samudrala Gourinath, Sankar Maiti

**Author notes:** Corresponding author: Sankar Maiti. Contribute equally (Priyanka Dutta, A.S. Jijumon, Mohit Mazumder).

## Abstract

Type VI secretion systems (T6SS) plays a crucial role in *Vibrio cholerae* mediated pathogenicity and predation. Tip of T6SS is homologous to gp27/gp5 complex or tail spike of T4 bacteriophage. VgrG-1 of *V. cholerae* T6SS is unusual among other VgrG because its effector domain is trans-located into the cytosol of eukaryotic cells with an additional actin cross-linking domain (ACD) at its C terminal end. ACD of VgrG-1 (VgrG-1-ACD) causes T6SS dependent host cell cytotoxicity through actin cytoskeleton disruption to prevent bacterial engulfment by macrophages. ACD mediated actin cross-linking promotes survival of the bacteria in the small intestine of humans, along with other virulence factors; establishes successful infection with the onset of diarrhoea in humans. Our studies demonstrated VgrG-1-ACD can bind to actin besides actin cross-linking activity. Computational analysis of ACD revealed the presence of WH2 domain through which it binds actin. Mutations in WH2 domain lead to loss of actin binding *in vitro*. VgrG-1-ACD having the mutated WH2 domain cannot cross-link actin efficiently *in vitro* and manifests less actin cytoskeleton disruption when transfected in HeLa cells.

**Summary statement:** Actin cross-linking (ACD) domain of VgrG-1 toxin of Type VI secretion in *Vibrio cholera* has WASP Homology domain 2 (WH2) domain. ACD interact with actin through WH2 domain, WH2 is essential for ACD mediated cross-linking and disruption of actin cytoskeleton in the host cell.

## Introduction

Cholera is life threatening water borne diarrheal disease caused by the gram negative bacterium *Vibrio cholerae*. The O1 and O139 *V. cholerae* serogroups produce primarily an enterotoxin called the CTX or cholera toxin. Cholera toxin is the main cause of diarrhoea [1, 2 and 3]. Other accessory toxins from *V. cholerae* including hemaglutinin/protease *(hapA)* or hemolysin *(hlyA)*, zot, Ace, VgrG-1, VopF/L are also involved in the pathogenesis [4 and 5]. Among these, Multifunctional Autoprocessing Repeats-in-Toxins (MARTX) and VgrG (Valine-Glycine Repeat Protein G) toxins are important categories of toxins having a signature Actin Cross-linking Domain (ACD) [6]. Type VI secretion systems (T6SS) is known to secrete three related proteins-VgrG-1, VgrG-2 and VgrG-3. VgrG components of the T6SS apparatus are capable to assemble and form a drilling device analogous to tail spikes in phages [7]. VgrG has similar structure resembling gp27/gp5 complex or the tail spike of T4 bacteriophage with an additional actin cross-linking domain at its C-terminal end that covalently cross-links the globular actin, leading to disruption of the host intrinsic actin cytoskeleton arrangements and cause rounded cell morphology [8 and 9]. The VgrG-1 of *V. cholerae* (T6SS) is unique compared to other VgrG, because its ACD containing effector domain is trans-located into the cytosol of eukaryotic cells [10]. In short, VgrG-1 helps *Vibrio* to prevent amoebic predation, to resist competing organisms for the colonization, to expand their niche and to infect host cells for the survival and propagation [11, 12, 13 and 14]. The VgrG-1-ACD is highly conserved, having a strong homology with the ACD of MARTX of *V. cholerae* [6].

Earlier studies have shown that VgrG-1-ACD is necessary for T6SS dependent host cell cytotoxicity and impairment of actin cytoskeleton mediated phagocytosis to prevent bacterial engulfment by macrophages [15 and 16]. The V-shaped three-dimensional structure of VgrG-1-ACD chemically cross-links two actin monomers through an isopeptide linkage. The bond is formed between 270^th^ glutamate amino acid residue of one actin monomer with 50^th^ lysine amino acid residue of another actin monomer. In spite of the structural details of the VgrG-1-ACD, the molecular mechanism of interaction of actin with ACD is poorly understood [9, 17 and 18].

In this study we have shown that presence of WH2 domain within the VgrG-1-ACD. The ACD binds to actin through this WH2 domain. Certain mutations in WH2 domain reduced the actin binding and actin cross-linking activity of ACD. The *in vitro* cell biology data also suggests that these mutations within the WH2 domain of ACD restrain the cross-linking activity of ACD. Consequently, we have shown that WH2 domain is essential for VgrG-1-ACD mediated actin filament disruption inside Hela cell.

## Results

### VgrG-1-ACD cross-links actin

Genomic DNA was isolated from *V. cholerae* O139 (Co 842) strain. The VgrG-1-ACD (732^nd^ - 1163^rd^) aa (Fig 1A) was cloned in pET-28a vector and expressed as N-terminal 6X-His tag protein and purified using Ni-NTA beads, further purified by gel filtration chromatography (Fig 1B). The cross-linking activity of the purified VgrG-1-ACD was examined by incubating 5 μM rabbit muscle actin (RMA) and different concentration of VgrG-1-ACD simultaneously in F-buffer (10 mM Tris pH 8.0, 0.2 mM ATP, 0.2 mM CaCl2, 0.2 mM DTT, 2.0 mM MgCl2 and 50 mM KCl) for 60 minutes. Subsequently the samples were checked in coomassie stained SDS-PAGE which showed the formation of oligomers of actin with higher molecular weight like dimer (~86 kDa), trimer (~128 kDa), tetramer (~170 kDa) and higher molecular weight oligomers. The formation of dimer, trimer and higher order oligomer of actin as an outcome of the cross-linking activity of ACD on actin (MW of monomer actin: 42.5 kDa) (Fig 1C) delineate the cross linking activity of VgrG-1-ACD from Co 842 is resembling with the VgrG-1-ACD of other strains of *V. cholerae* [19].

**Figure 1:**
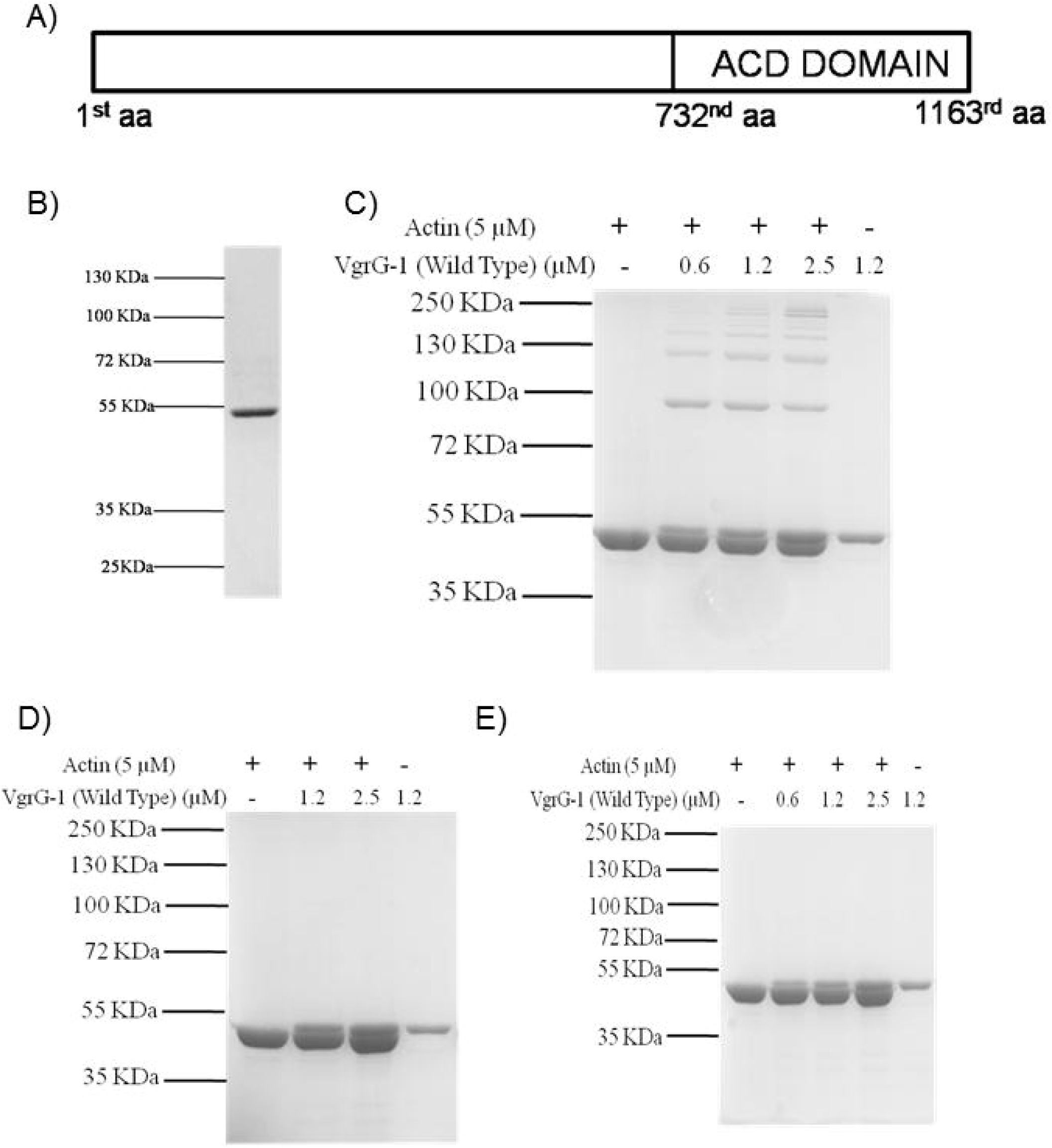
Valine-Glycine Repeat Protein G (VgrG-1), having ACD (Actin Crosslinking Domain) with actin cross-linking activity. **[A]** Schematic representation of *V. cholerae* VgrG-1 toxin domain showing ACD from (732^nd^ aa-1163^rd^ aa). **[B]** Purified 6X His tag full length VgrG-1-ACDprotein (50KDa) in a coomassie stained 12% SDS-PAGE. **[C]** Wild type VgrG-1-ACD protein was found to cross-link actin in a concentration dependent manner (N=6). **[D]** In vitro Cross-linking activity of VgrG-1-ACD protein incubated with preformed F-actin (N=4). **[E]** In vitro cross-linking assay of VgrG-1-ACD protein incubated with G-actin in G buffer (N=4).

### VgrG-1 ACD fails to cross-link F-actin and G-actin

We incubated 5 μM RMA in F-buffer for 45 minutes to form F actin prior to addition of VgrG-1-ACD. The ACD was added in the reaction mixture of F-actin and kept for another 15 minutes. SDS-PAGE result demonstrated that absence of cross-linked actin with higher order molecular weight other than 42.5 kDa band of actin monomer (Fig 1D). This illustrated that the addition of VgrG-1-ACD after the formation of F-actin failed to cross-link F-actin (Fig 1D).

Further we liked to test the cross-linking activity of VgrG-1-ACD with G-actin. We had simultaneously incubated 5 μM RMA with VgrG-1-ACD in G-buffer (10 mM Tris pH 8.0, 0.2 mM ATP, 0.2 mM CaCl_2_, and 0.2 mM DTT) for 60 minutes. Our SDS-PAGE results depictedthe presence of monomeric actin band with molecular weight 42.5 kDa and absence of oligomers of actin bands with higher molecular weight (Fig 1E). This result exhibited that VgrG-1-ACD was also insufficient to cross-link monomeric G-actin (Fig 1E) as shown previously [20].

### VgrG-1-ACD co-sediments with F-actin

We were curious to study the mechanism of interaction of VgrG-1-ACD with actin. Furthermore it was investigated by addition of 5 μM RMA withVgrG-1-ACD in a concentration dependent manner simultaneously in F-buffer and reaction mixtures were incubated for 60 minutes followed by co-sedimentation using high speed centrifugation at 310X100 g for 45 minutes. The supernatant and pellet fraction were separated out in SDS PAGE. We observed that VgrG-1-ACD had cross-linked with actin in the supernatant fractions forming higher molecular weight oligomers. The pellet fractions in the SDS PAGE illustrated that it was bound to the filamentous actin in a concentration dependent manner (Fig 2). In case of VgrG-1-ACD control, in which there was no actin, the protein was only in the supernatant fraction. This indicated that to perform the cross-linking reaction, VgrG-1-ACD should bind to actin. The same reaction conditions were followed except VgrG-1-ACD was added after 45 minutes of formation of F-actin, surprisingly we can see that VgrG-1-ACD bound to F-actin in the pellet fraction but no cross-linking activity in the supernatant fraction was observed even in higher concentration of VgrG-1-ACD (Fig S1A). Remarkably these results confirmed that VgrG-1-ACD has actin binding ability.

**Figure 2:**
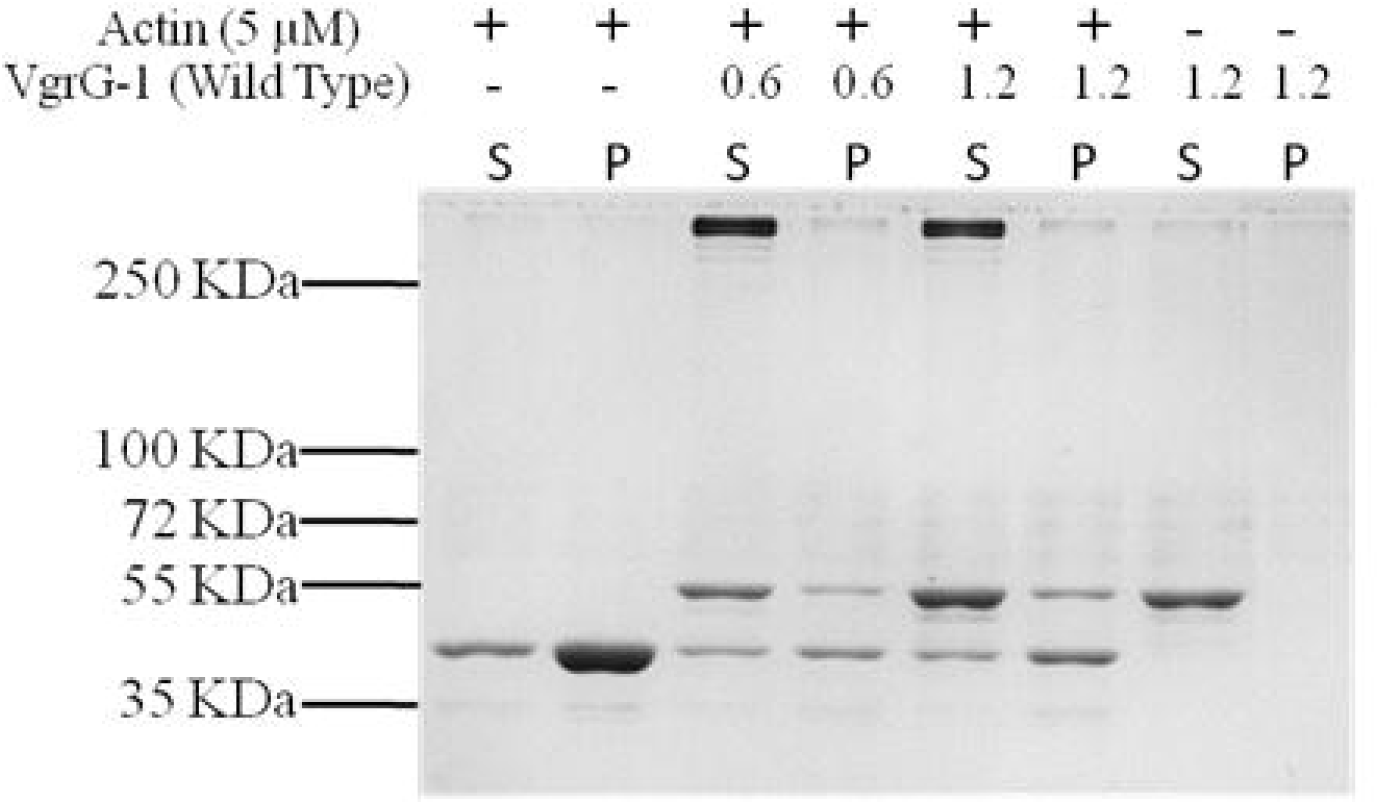
Co-sedimentation of VgrG-1-ACD of *V. cholerae* with F-actin by cosedimentation assay. S and P indicate the supernatant and pellet respectively. The concentration of actin was 5μM. VgrG-1-ACD was added in an increasing concentration. RMA and VgrG-1-ACDwere added simultaneously in the reaction mixture and centrifuged at 310 x 1000 g for 45 minutes (N=3).

### Presence of putative WH2 domain in VgrG-1-ACD

To quest the actin binding region in VgrG-1-ACD, we performed profile based search using the protein sequence of VgrG-1-ACD. The crystal structure of VgrG-1-ACD of *V. cholerae* which was already reported [9] along with WH2 domains from higher organisms like *Homo sapiens, Mus musculus* and *Bos taurus* were considered for analysis. The result of these analysis suggested that WH2 domain in VgrG-1 was poorly conserved which was a common characteristic of the WH2 domains and the alignment clearly showed that the probable WH2 domain was present around (1009^th^ - 1020^th^) aa (Fig 3A) [21]. The sequence similarity based tree generated using Clustal Omega [22] (Fig 3B) with the known WH2 domains demonstrated high sequence diversity in WH2 domain even amongst higher eukaryotes. The actin binding WH2 domain was identified on the basis of sequence conservation, evolutionary function and structure of the domain. The probable WH2 domain was located on the surface of the ACD (Fig 3C) with the conserved residue Ile-1017 placed in the middle of the 6^th^ alpha helix (Fig 3C’) in the crystal structure. The hotspot analysis using KFC server [23] for WH2 domain indicated Val-1013, Met-1014, Ile-1017 and Lys-1018 as binding hotspots. The (Fig 3C) depicted the location of hotspots at the binding interface of G-actin. To further validate the presence of WH2 domain, we extracted all the structures of WH2 domains co-crystallized with actin monomer and in some cases actin-DNase-WH2 complex. The WH2 domains were further extracted, superimposed (Fig 3D and E) and used for structure based sequence alignment, which clearly showed that the residues in the ACD domain (1009^th^-1020^th^) fold in a similar way and has the conserved residues that bind to actin (Fig 3D and E) in the experimental structures. Our sequence analysis displayed that VgrG-1-ACD amino acids (1009^th^-1020^th^) “DAKAVMKHIKPQ” aligned appropriately with other known WH2 sequences (Fig 3A). The (Fig 3C) indicated the location of hotspots at the binding interface of G-actin which was need to be tested further using in vitro reconstitution assays.

**Figure 3:**
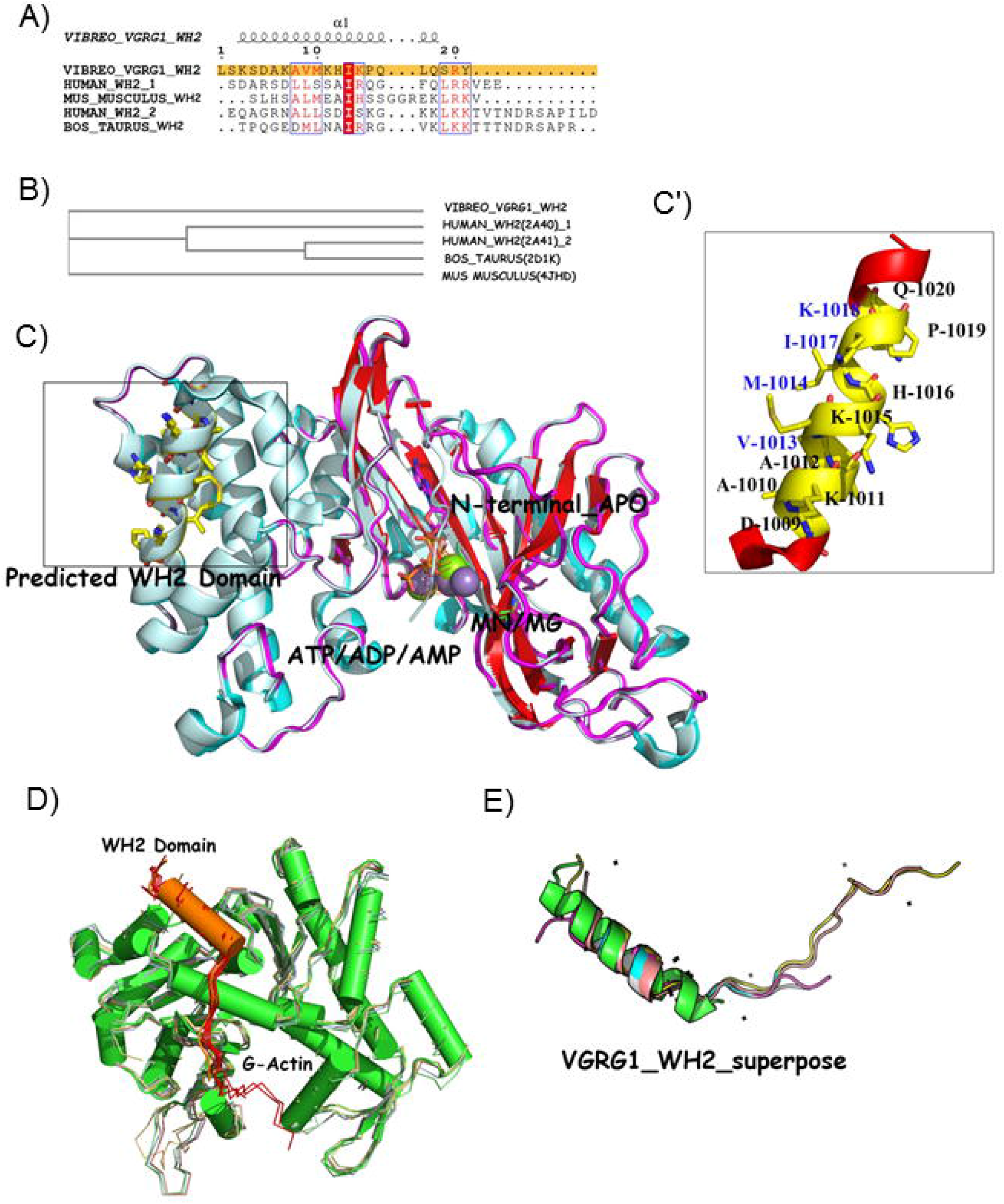
Structural and sequence analysis of VgrG-1 actin binding WH2 domain within the ACD. **[A]** Structural alignment of ACD WH2 domain with known WH2 domain sequences taken from co-crystallized structures of actin. The sequence of ACD-WH2 domain of VgrG-1 shared significant identity with proteins having a characterized WH2 domain. The sequences of WH2 domain extracted and aligned from the actin complexes of WASP and WAVE domains from Human (2A40 & 2A41), actin-WH2 domain from *Bos taurus* (2D1K) and *Mus musculus* (4JHD). **[B]** The phylogenetic tree derived from the alignment showing ACD-WH2 domain of VgrG-1 and *Mus musculus* WH2 diverged into different branches whereas the WH2 domains from WASP, WASP-family verprolin homologous protein from *Homo sapiens* and *Bos taurus* WH2 domain are branched together. **[C]** The superimposition of VgrG-1-ACD co-crystalized with AMP, ADP and ATP with two divalent metals bound at the substrate binding site showed a V-shaped structure, comprising of nine α helices and twelve β strands. [C’] The zoomed view of the predicted structure of WH2 like domain showed its presence at the surface of the ACD domain as the 6^th^ alpha helix structure with amino acid sequence “DAKAVMKHIKPQ”. **[D]** and **[E]** Structural superimposition of all the structure of WH2 domain co-crystallized with actin along with the probable ACD WH2 domain of VgrG-1.

### Predicted WH2 domain within VgrG-1-ACD binds to actin but do not cross link actin

Based on the above bioinformatics analysis of WH2 domain within the ACD, we had sub-cloned the WH2 domain (998^th^-1099^th^) aa of ACD (ACD-WH2) (Fig 4A) in pET-28a vector and purified as N-terminal 6X-His tagged protein using Ni-NTA beads and further purified by gel filtration chromatography (Fig 4B). Different concentration of ACD-WH2 was incubated for 60 minutes with 5 μM of RMA in F-Buffer and further high speed centrifugation was done as above mentioned. SDS PAGE analysis showed ACD-WH2 domain had co-sedimented along with actin in the pellet fraction in a concentration dependent manner. Simultaneously, in the control the WH2 domain in absence of actin showed up in the supernatant fraction only. This indicated that ACD-WH2 was efficient to bind actin (Fig 4C). As WH2 domain is an ancient actin binding motif found in eukaryotes regulating actin cytoskeleton [24 and 25] so to validate the WH2 interaction with actin, nucleation assay was performed with 4 uM RMA mixed with pyrene-RMA and ACD-WH2. Actin nucleation was inhibited by ACD-WH2 in a concentration dependent manner (Fig 4D). This might be possible only if ACD-WH2 able to sequestered actin monomer. Therefore ACD-WH2 was able to successfully interact with actin (Fig 4C and 4D) [26]. The dissociation equilibrium constant (K_D_) was approximately 50 nM which was in the range of other WH2 domain which binds to actin (Fig 4G) [27 and 28]. The WH2 domain within VgrG-1-ACD was not able to cross link actin like the VgrG-1-ACD (Fig S1C), which illustrated that ACD-WH2 domain was necessary for efficient substrate binding for proper cross-linking activity of V grG-1 - ACD.

**Figure 4:**
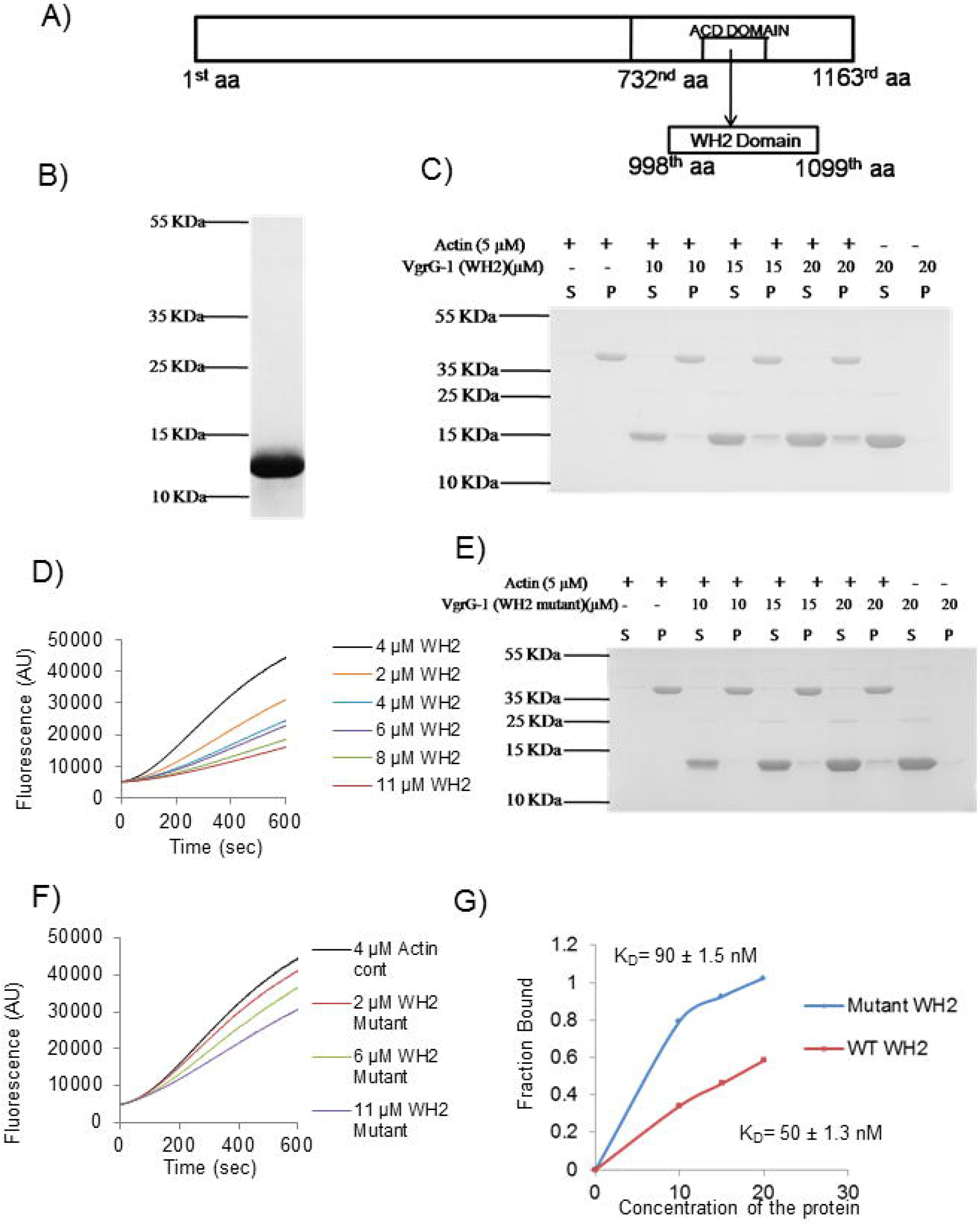
Proposed WH2 domain within ACD was able to bind F-actin in co-sedimentation assay. **[A]** Proposed domain structure of WH2 like domain in the VgrG-1-ACD (998^th^ aa – 1098^th^ aa). **[B]** The commassie stained 15% SDS PAGE showed purified protein of WH2 like domain (11KDa). **[C]** The co-sedimentation assay showed WH2 domain could bind actin in a concentration dependent manner (N=4). **[D]** Monomer sequestering assay of WH2domain using 10% pyrene labelled actin (N=4). **[E]** The co-sedimentation assay of mutant WH2 domain (N=4). **[F]** Monomer sequestering assay of mutant WH2 domain of VgrG-1 using 10% pyrene labelled actin (N=4). **[G]** The binding affinity of wild type and mutant WH2 domain to F-actin.

### Reduction in actin binding ability with mutation in the WH2 domain in VgrG-1-ACD

Amino acids from the hot spots of actin binding site of ACD-WH2 domain, predicted from bioinformatics analysis (Fig 3) were V (1013), M (1014), I (1017) and K (1018) [23 and 29]. We mutated these amino acids to alanine following the site directed mutagenesis protocol. The mutated WH2 domain within ACD (ACD-mWH2) containing these mutations V1013A, M1014A, I1017A and K1018A was purified as N-terminal 6X-His tagged, using Ni-NTA beads. Different concentration of ACD-mWH2 was mixed with actin and F-Buffer and co-sedimented using high speed similarly as mentioned above. The SDS PAGE data showed that ACD-mWH2 co-sedimented to lesser extent in comparison to the wild type (Fig 4E). The fluorometric 4 μM nucleation assay showed that the ACD-mWH2 was unable to sequester actin monomers as efficiently as the wild type (Fig 4F). Therefore this indicated that ACD-mWH2 was inefficient to interact with actin. Our co-sedimentation assay data exhibited the binding affinity of ACD-WH2 to actin was more than that of the ACD-mWH2 (Fig 4G).

### The VgrG-1-ACD containing mutated WH2 domain cannot cross-link actin

We found that ACD-WH2 was able to interact with actin efficiently than ACD-mWH2 construct. These results suggested that the WH2 domain present within the ACD region facilitated efficient communication of ACD with actin. This made us curious to investigate the consequence of the actin cross-linking activity of VgrG-1-ACD containing the mutated WH2 domain. ACD containing I1017A and K1018A WH2 mutation was purified as N-terminal 6X-His tagged, using Ni-NTA beads (Fig S1B). Actin cross-linking experiments following above protocol was performed and the data showed that IK to AA mutation in WH2 domain of ACD led to less actin oligomers formation as compared to that of wild type VgrG-1-ACD (Fig 5A). Following this, VgrG-1-ACD containing V1013A and M1014A WH2 mutation was purified as above (Fig S1D) and actin cross-linking assay was performed. The results depicted that VM to AA mutation in WH2 domain of ACD resulted in loss of actin cross-linking activity of ACD (Fig 5B). Next we had prepared a construct of VgrG-1-ACD containing V1013A, M1014A, I1017A and K1018A WH2 mutation and the construct was purified as above (Fig S1E) and cross-linking assay was performed. The results explicitly implied that VgrG-1-ACD containing V1013A, M1014A, I1017A and K1018A WH2 mutation was incompetent to cross-link actin and forming higher order actin oligomers (Fig 5C). Previously we had seen that the WH2 mutant with the above mutations did not communicate with actin efficiently (Fig 4E and 4F) suggesting the necessity of WH2 domain for effective actin interaction and subsequent cross-linking activity of VgrG-1-ACD.

**Figure 5:**
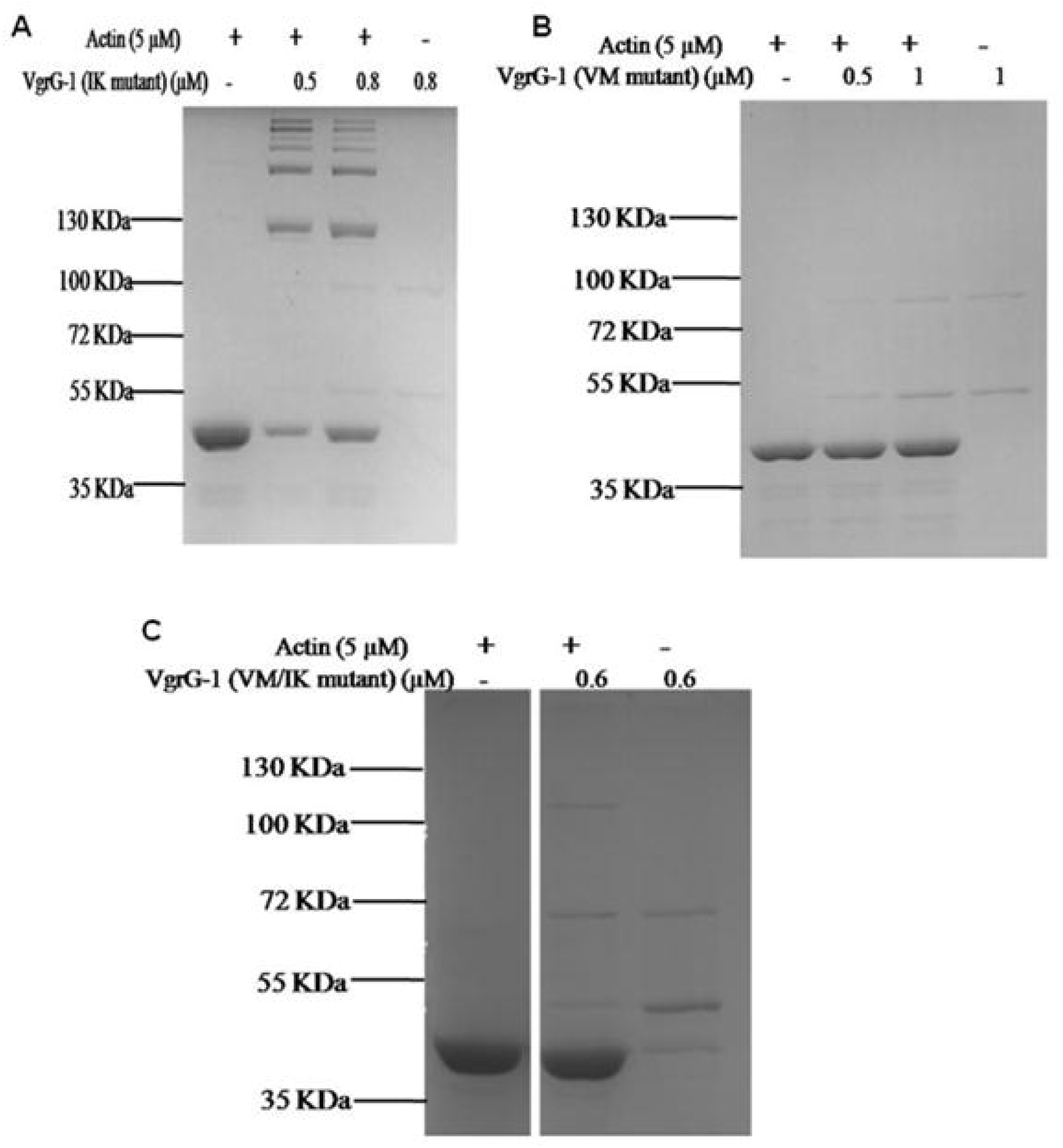
VgrG-1-ACD of the *V. cholerae* having mutations V1013A & M1014A and I1017A & K1018A was not able to cross-link actin *in vitro* comparing to the wild type. **[A]** Cross-linking activity of VgrG-1ACD having mutations I1017A & K1018A. **[B]** Cross-linking assay of V1013A and M1014A mutant protein showing no cross-linking activity. **[C]** Cross-linking activity of VgrG-1-ACD domain having mutations I1017A & K1018A and V1013A & M1014A showing no cross-linking activity (N=4).

### The VgrG-1-ACD destroys the actin cytoskeleton but mutation leads to loss of crosslinking activity in HeLa cells

Later on we evaluated the role of actin cross-linking activity by VgrG-1-ACD in *in vitro* cell biology using cultured HeLa cells. The full length wild type VgrG-1-ACD was sub-cloned in pEGFP-C1 vector as C-terminal GFP tagged and transfected in HeLa cells and stained with phalloidin after 36 hours to study the actin cytoskeleton. The transfection experiment exhibited that the full length VgrG-1-ACD disrupted the actin cytoskeleton (Fig 6B). But in case of control pEGFP-C1 transfected HeLa cells, normal intact actin fibres were observed (Fig 6A). Then we had sub cloned ACD-WH2 in pEGFP-C1 vector as N-terminal GFP tagged and transfected in HeLa cells and stained with phalloidin to see the actin cytoskeleton. Results showed that only WH2 domain did not disrupt actin cytoskeleton (Fig 6C) which strengthen our previous data that only WH2 domain did not cross link the actin (Fig S1C). Finally we had sub cloned VgrG-1-ACD containing V1013A, M1014A, I1017A and K1018A WH2 mutation in pEGFP-C1 vector as N-terminal GFP tagged and transfected in HeLa cells and stained with phalloidin to examine the actin cytoskeleton. Transfection results demonstrated that the VgrG-1-ACD mutant with both VM to AA and IK to AA did not able to disrupt actin fibres, actin fibres were similar that of control (Fig 6D). This confirmed that WH2 domain mediated actin binding was also essential and imperative for efficient actin cross-linking activity of VgrG-1-ACD.

**Figure 6:**
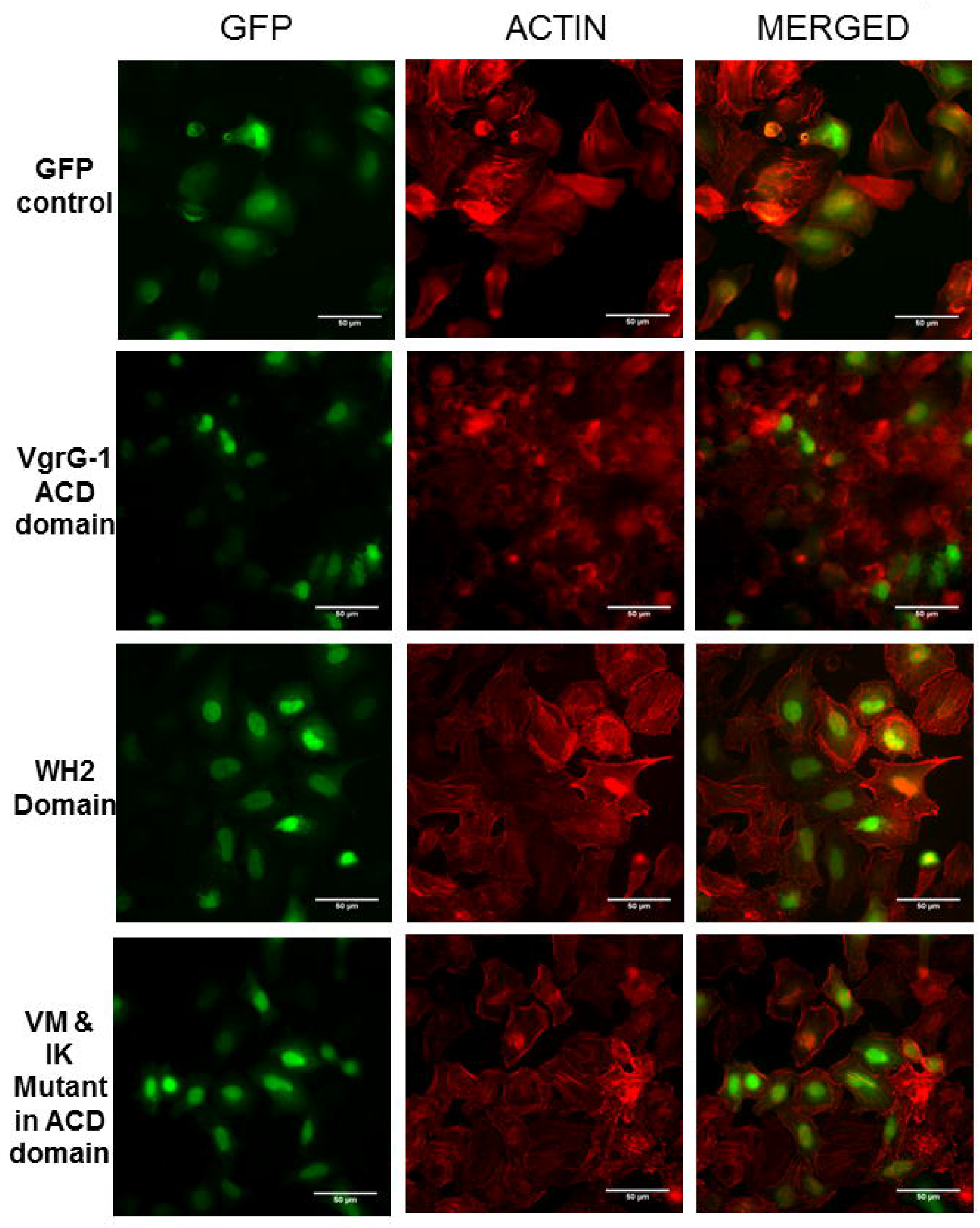
Effect of VgrG-1-ACD and WH2 fragments in cellular actin cytoskeleton. **[A]** The GFP transfected HeLa cells showed no particular change in rhodamine phalloidin stained actin cytoskeleton showed in first row second column. [B] HeLa cells transfected with GFP tagged full length VgrG-1-ACD domain (732^nd^ aa-1163^rd^ aa) showed disrupted rhodamine phalloidin stained actin cytoskeleton second row third column. [C] HeLa cells transfected with GFP tagged WH2 like domain fragment did not show any change in rhodamine phalloidin stained actin cytoskeleton second column third row. [D] The ACD domain mutant (V1013A & M1014A and I1017A & K1018A) tagged with GFP did not able to disrupt the actin cytoskeleton. The expression of GFP tagged vectors is shown in first column, the rhodamine phalloidin stained actin cytoskeleton is shown in second column and the merged of the two were shown in third column. All images were captured and scaled with identical settings. The scale bar is 50 μm. The magnification is 60X (N=5).

### Computational model depicts favourable binding of VgrG-1-ACD via WH2 domain with actin

With the knowledge that the identified WH2 domain bound with actin, we looked for possible modes of interaction of VgrG-1-ACD with G and F-actin using structural superimposition and Cluspro docking software. Based on the structural superimposition here we showed the possible orientation or possible mode of interaction of the 6^th^ helix that was present in the WH2 domain of VgrG-1. The WH2 domain modelled with G–actin on the basis of structural similarity, it showed that it was able to bind between subdomains 1 and 3 (Fig 7A). The protein–protein blind docking simulations were successfully carried out using three adjacent actin monomers taken from the filamentous actin structure. In blind docking simulations the active site was not defined for any of the proteins, hence the VgrG-1-ACD was allowed to bind in all possible actin cavities, after docking the top docking poses from a cluster of 37 structures were used for the surface representation in (Fig 7B). The docking results manifested that the C-terminal of the VgrG-1 alpha helix containing the WH2 domain was bound in a head on manner with the filamentous actin. The ACD-WH2 domain was binding between the subdomain 1 and subdomain 3 (Fig 7C) of actin-2 and the N-terminal part of the ACD domain interacts with actin-1. The K-50 residue proposed to be involved in the cross-linking was present at the binding interface of subdomain1 and subdomain 2 where the WH2 domain was bound to actin, whereas the E-270 was present at the binding interface of the actin filaments (Fig 7D). The docking studies and the superimpose model clearly suggested that the WH2 domain was capable of binding both G-actin and F-actin. The docked domain (three actin molecule andVgrG-1-ACD) was then energy minimized and used to build a full length model of filamentous actin using structural superimposition.

**Figure 7:**
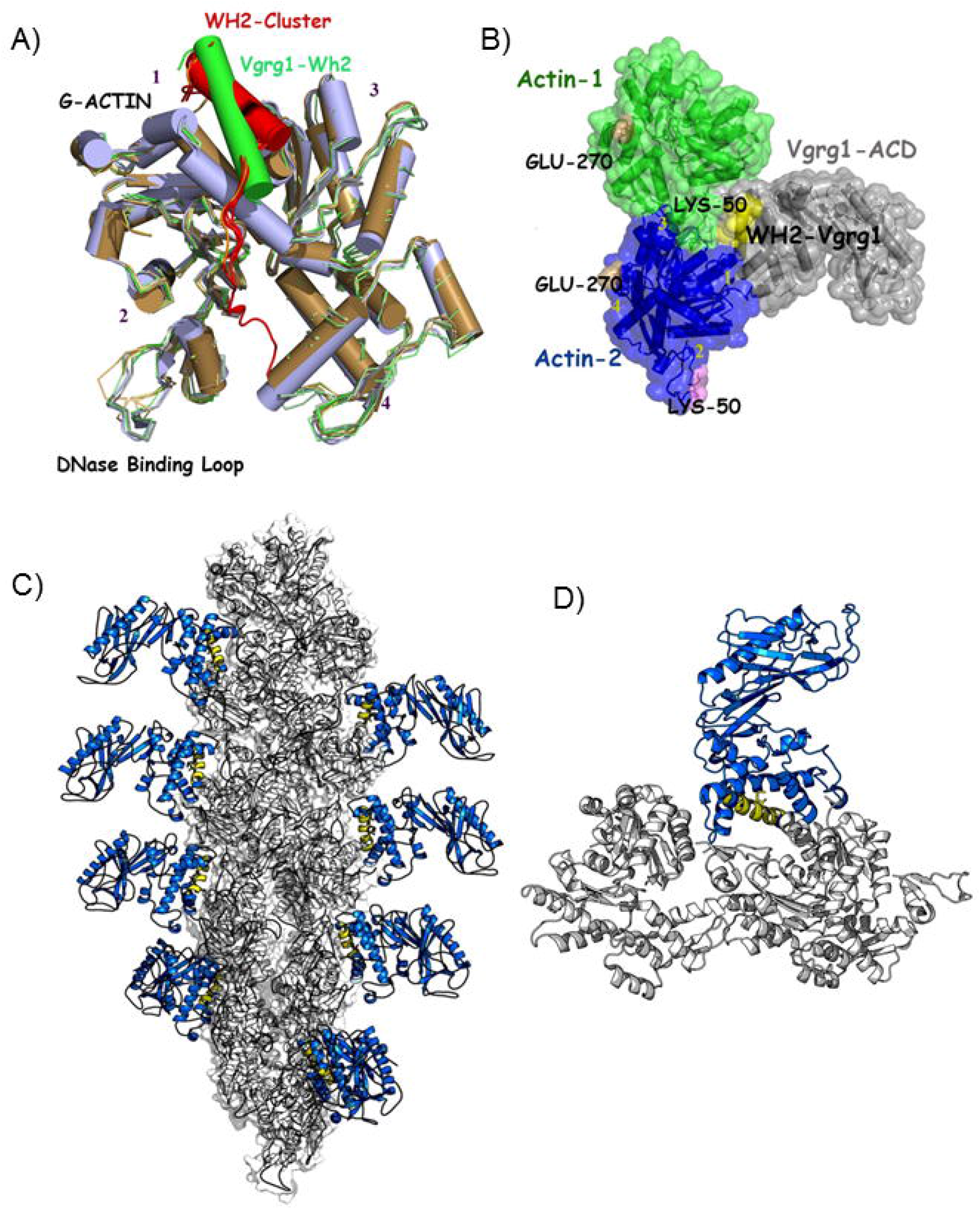
Proposed molecular modeling of actin binding of WH2 of ACD domain of VgrG-1 based on molecular docking and structural superposition. **[A]** The model showed the superposed cluster of WH2 domain crystallized with actin from different organisms along with WH2 domain of VgrG-1. **[B]** The docked model of filamentous actin showing two actin monomers interacting with the WH2 of VgrG-1 which was bound between the subdomain1 and subdomain3 and was in close proximity with K50 crosslinking residue in actin-1. **[C]** The F-actin decorated ACD domain complex was obtained by using superimposition of initial model obtained from Cluspro into the different chains (Monomers) in F-actin complex. **[D]** The actin has been shown in grey colour, ACD domain in blue and the binding WH2 domain is shown in yellow.

## Discussion

More than 150 actin binding proteins are known to regulate actin filament dynamics [30 and 31]. VgrG-1-ACD had been discovered as actin cross-linking protein which forms non polymerizable actin oligomers. We found that the VgrG-1-ACD from O139 strain (Fig 1C) was able to cross link actin very efficiently similar to VgrG-1 of other serogroups of *Vibrio cholerae* [20]. We had seen that VgrG-1-ACD and RMA incubated simultaneously in F-buffer resulted in efficient actin cross-linking activity of VgrG-1-ACD (Fig 1C). When VgrG-1-ACD was added to the pre formed F-actin, the capability to cross link actin was completely diminished. This indicated that F-actin was not a good substrate for the VgrG-1-ACD (Fig 1D). On the other hand, VgrG-1-ACD incubated with G-actin in G-buffer resulted in no actin cross-linking (Fig 1E) ruling out that G actin was also not a good substrate for the VgrG-1-ACD similar to earlier results showing _+2_ that Mg^+2^ was necessary for the cross linking activity [20].

VgrG-1-ACD catalysed the covalent bond formation between two actin monomers [17 and 18]. For this activity, VgrG-1-ACD must first bind to actin. Our results revealed that VgrG-1-ACD was able to co-sediment with actin in a concentration dependent manner; when actin, VgrG-1-ACD and F-buffer was added simultaneously (Fig 2) or VgrG-1-ACD was added 45 minutes after the formation of F-actin (Fig S1A). In the latter case VgrG-1-ACD was unable to cross-link actin even in presence of high concentration of VgrG-1-ACD because nascent actin oligomers are the appropriate substrate for crosslinking activity of VgrG-1-ACD.

The above results validate that VgrG-1-ACD could bind actin. Hence actin binding domains might exist within VgrG-1-ACD. Further computational analysis illustrated that VgrG-1-ACD contains a putative WH2 domain (Fig 3). WH2 domain is well characterized actin binding domain [21 and 25]. The presence of WH2 domain in *Vibrio* is not a surprise as *Vibrio* has other effector proteins like VopL, VopF which contain multiple WH2 domains and modulate the infected host cell actin dynamics [28, 32 and 33]. The presence ACD-WH2 domain was demonstrated with aid of sequence homology, structural and evolutionary functional significance of the domain. ACD-WH2 domain folds exactly the same as seen in other WH2 co-crystallized with G-actin (Fig 3C) [21], suggesting that the identified domain might work similarly. The hotspot analysis was done using KFC server for WH2 domain indicating Val-1013, Met-1014, Ile-1017 and Lys-1018 as the crucial amino acids of the binding hotspots.

For further confirmation of actin binding ability of ACD-WH2, we had co-sedimented the purified predicted ACD-WH2 with actin using high speed centrifugation. The assay revealed that ACD-WH2 co-sedimented with actin in a concentration dependent manner (Fig 4C) indicating that the predicted ACD-WH2 could bind actin. Additionally the inhibition of 4 μM actin nucleation by the ACD-WH2 supported its presence within VgrG-1-ACD (Fig 4D), as it is established that WH2 domain interfere with the 4 μM fluorometric actin nucleation assay by sequestering actin monomers [25]. The ACD-mWH2 co-sedimented with actin in lesser amount as compared to wild type and also not able to inhibit actin nucleation efficiently like that of wild type (Fig 4E and 4F). This illustrated that VgrG-1-ACD interacts with actin through its WH2 domain. Results of actin cross-linking assay with VgrG-1-ACD having WH2 mutation showed less or no actin cross-linking. The IK to AA mutant cross-link actin less efficiently than the wild type but the VM to AA mutant did not able to cross link actin and the fragment containing both VM and IK mutated to AA rendered no cross-linking of actin (Fig 5). This confirmed that the interaction of VgrG-1-ACD with actin through its WH2 domain is also important for actin cross-linking activity. This result was further validated by *in vitro* cell biological data of phalloidin stained HeLa cells transfected with VgrG-1-ACD and ACD with VM and IK mutated to AA in WH2 domain. Results showed only the VgrG-1-ACD is able to disrupt the actin fibres inside the HeLa cells but neither the ACD containing mutated WH2 domain nor the ACD-WH2 alone can disrupt the actin cytoskeleton (Fig 6) [34]. The cell data affirmed with the *in vitro* actin reconstitution assays (Fig 1C, 5C, S1A and S1C); all together suggesting that VgrG-1-ACD binds actin through its WH2 domain and this actin binding is also indispensable for ACD mediated actin cross-linking along with previously studied mutations [9].

Finally, our model showed that (Fig 7) VgrG-1-ACD was able to interact with actin at the interface of two actin monomer through its WH2 domain. Based on the proposed model of F-actin-ACD complex, we could claim that the WH2 domain binds to two actin monomers (Fig 7) where WH2 helix interacts with domain I of one actin monomer and domain II of another actin which is similar to earlier reports, the chemical cross-linking between two different actin monomers [17]. In G-actin, WH2 helix enters in the groove between domain 1 & 3. This complex model obtained using docking simulation that is proposed in this paper gives insight to the mode of binding of ACD with F-actin [35]. The proposed model was devised on the basis of biochemical data reported in this study, especially the properties of the mutants.

VgrG-1-ACD toxin had been reported to forms non polymerizing non-functional actin oligomers from monomeric actin to cause successful pathogenesis [36 and 37]. Our data suggest that VgrG-1-ACD binds to F-actin through WH2 domain. In future, co-crystal structure of actin and VgrG-1-ACD would illuminate light on the exact molecular mechanism of actin crosslinking mediated byVgrG-1-ACD.

## Material and methods

### Plasmid constructs

*V. cholerae* genomic DNA was isolated from O139 (C_0_842) strain. We amplified the construct of VgrG-1-ACD (732^nd^ – 1163^rd^aa) by PCR from the genomic DNA of *V. cholerae* using the forward primer 5′ GGAAGATCTCCAACACATTTCCCGAAGTC 3′ and reverse primer 5′ CGCAAGCTTTTATTGCCATTCTTGAGGATTATGC 3′ with Bgl II and Not I restriction enzyme sites respectively. The PCR product was cloned in pET-28a vector (Novagen). In a similar way the constructs of WH2 domain were sub cloned from the VgrG-1-ACD of *V. cholerae* using forward primer 5′ GGAAGATCTCCAAGCAAAATAGTAGAAATACTC 3′ and reverse primer 5′ CGCAAGCTTTTACTTCATAGGATCAAACTGATCG 3′ in pET-28a vector with BglII and HindIII restriction enzyme sites respectively.

### Site directed mutagenesis

The VgrG-1-ACD clone in pET-28a was used as template, with forward and reverse primers having desired mutations, amplified the entire plasmid. The PCR was carried out using Pfu turbo DNA polymerase (Stratagene) having a polymerization rate of 1Kb/min. The non-mutated parental amplicons were digested with DpnI and the remaining amplicons were transformed into DH5α *E.coli* competent cells. We confirmed the mutations by DNA sequencing from the screened mutant clones.

### I1017A and K1018A

Forward primer:

5′ CCAAAGCGGTTATGAAACAT**GCCGCG**CCTCAATTGCAGTCTCGC3′

Reverse primer:

5′ GCGAGACTGCAATTGAGGCGCGGCATGTTTCATAACCGCTTTGG3′

### V1013A and M1014A

Forward primer:

5′ GAGTGATGCCAAAGCG**GCCGCG**AAACATATCAAGCCTCAATTG3′

Reverse primer:

5′ CAATTGAGGCTTGATATGTTTCGCGGCCGCTTTGGCATCACTC3′

### I1017A, K1018A, V1013A and M1014A

Forward primer:

5′ AAGAGTGATGCCAAAGCG**GCCGCG**AAACAT**GCCGCG**CCTC3′

Reverse primer:

5′ GAGGCGCGGCATGTTTCGCGGCCGCTTTGGCATCACTCTT3′

### Protein purification

G-actin was purified from rabbit skeletal muscle acetone powder [38]. Actin was labelled with N-(1-pyrene) iodoacetamide (P-29, Molecular Probes) [39]. The unlabelled and labelled actin was purified using Hiprep 16/60 Sephacryl S-200 gel filtration column (GE Healthcare). The column was equilibrated with G-buffer (2 mM Tris pH 8.0, 0.2 mM ATP, 0.2 mM CaCl_2_ and 0.2 mM DTT).

For purification of the proteins, the expression constructs were transformed into *E. coli* BL21 DE3 (Stratagene) cells. The construct containing *E. coli* cultures were grown up to O.D 0.6-0.8 in rich media (20 gm Tryptone/liter, 10 gm Yeast Extract/liter, 0.58 gm NaCl/liter, and 0.18 gm KCl/liter) with 30μg/ml Kanamycin (USB) at 37°C. When the O.D_600_ crossed 0.6 the temperature was kept 37°C and induced with 0.5 mM IPTG (isopropyl-1-thio-β-D-galactopyranoside, Fermentas) and subsequently the cultures were grown for 4 hours and pelleted down at 6000 rpm for 15 minutes and stored in −80°C freezer.

We did all the steps of protein purification at 4°C or keeping them on ice [40]. Pelleted bacterial cells were taken and resuspended in lysis buffer (50 mM Tris-Cl pH 8.0, 100 mM NaCl, 30 mM Imidazole pH 8.0, 1 mM DTT(USB), IGEPAL(Sigma-aldrich), Protease inhibitor cocktail(1mM Phenylmethylsulfonyl fluoride, Benzamidine hydrochloride, Leupeptin, Aprotinin, PepstatinA) and extracted by sonication. Then it was centrifuged at 12000 rpm for 15 minutes. The supernatant was incubated with 50% Ni-NTA slurry of the beads (Qiagen) for 2 hours. After that it was washed thrice with wash buffer (50 mM Tris-Cl pH 8.0, 20 mM NaCl, 30 mM Imidazole pH 8.0). The protein was eluted with elution buffer containing high concentration of Imidazole (50 mM Tris-Cl pH 8.0, 20 mM NaCl, 250 mM Imidazole pH 8.0 and 5% glycerol). Later on, the purified protein was dialyzed in HEKG_5_ buffer (20 mM HEPES, 1 mM EDTA, 50 mM KCl and 5% Glycerol) for overnight and further stored in 20 mM HEPES, 1 mM EDTA, 50 mM KCl and 10% Glycerol.

### Actin cross-linking assay

5 μM RMA was taken along with different concentration of VgrG-1-ACD and incubated for 1 hour at 25°C in F-buffer (10 mM Tris-Cl pH 8.0, 0.2 mM DTT, 0.7 mM ATP, 50 mM KCl, 2 mM MgCl_2_). The net volume of the reaction was 50μl and each set had one actin and protein control reaction mix. After completion of incubation, the samples were boiled with sample loading buffer and loaded in SDS-PAGE.

### High speed pelleting assay

Actin and different concentration of VgrG-1-ACD and ACD-WH2 domain was incubated at 25°C in F-buffer (10 mM Tris-Cl pH 8.0, 0.2 mM DTT, 0.7 mM ATP, 50 mM KCl, 2 mM MgCl_2_) for one hour. We set the net volume of the reaction to 50μl keeping the concentration of actin to 5μM in each reaction mix. As a control, we performed one reaction without adding protein and one reaction without adding actin. After completion of incubation, the samples were centrifuged at 310 x 1000 g for 45 minutes in TLA-100 rotor (Beckman Coulter). Subsequently, we separated the supernatant to a fresh tube and mixed it with SDS-PAGE sample loading buffer. Thereafter the removal of the supernatant, the pellet was resuspended in F-buffer and SDS-PAGE sample loading buffer was added. Then the supernatants and pellets were analysed in 12% coomassie-stained SDS-PAGE and K_D_ was calculated accordingly [41].

### Bioinformatics analysis of VgrG-1

One of the salient features of WH2 domain was; it short in length and poorly conserved [21]. Based on structural model [9]; we performed sequence based structural alignment of VgrG-1ACD with all the available crystal structures of WH2 domain. The identification of WH2 domain within the VgrG-1-ACD was carried out using HMM based model obtained from PFAM database using HMMER [42]. We further tried to align the predicted WH2 with the known crystal structures co-crystalized with actin monomer and in some cases actin-DNase-WH2 complex. The sequence and structural alignments were carried out using Clustal Omega [22], ESpript [43] and Promals3D server [44]. The identified sequence which folds into a helix as seen in the crystal structure was then subjected to hotspot prediction using KFC server [23 and 29] which predicted the binding “hot spots” within protein-protein interfaces by recognizing structural features indicative of important binding contacts.

### 4μM nucleation assay

Monomeric actin was labelled with N-(1-pyrene) iodoacetamide (P-29, Molecular Probes) [39]. Labelled and unlabelled actin was mixed in 1:9 ratio to prepare final stock concentration of 12 μM in G-buffer (10 mM Tris pH 8.0, 0.2 mM ATP, 0.2 mM DTT, 0.2 mM CaCl_2_). From this stock solution, required amount of actin was taken such that final concentration of actin is 4 μM in 60 μl reaction volume. Exchange buffer (10 mM EGTA, 1 mM MgCl2), 20X Initiation Mix (1 M KCl, 40 mM MgCl2, and 10 mM ATP) and candidate protein in different concentrations were added and the volume was adjusted with G-buffer and HEKG_5_. The reaction was excited at 365 nm and the emitted spectrum was recorded at 407 nm at 25°C in fluorescence spectrophotometer (QM40, Photon Technology International, Lawrenceville, NJ).

### Cellular experiments

HeLa cells (ATCC^®^ CRM-CCL-2™) were maintained in high glucose Dulbercco’s Modified Eagle’s Media (Gibco) containing 2 mM L-Glutamine (Gibco) Pen-Strep (Hyclone, Thermo Scientific) and 10% Fetal Bovine Serum (Gibco). We carried out the transfection experiments in 24-well plate using Lipofectamine 2000 (Invitrogen) following the manufacturer’s protocol. For this the cells were grown on poly D-lysine coated cover slips placed in wells. The cells were fixed using 4% paraformaldehyde (Sigma Aldrich) dissolved in PBS (phosphate buffer saline pH 7.4) for 30 minutes in room temperature after 36 hours of transfection. Subsequently the cells were washed thrice in PBS and the actin cytoskeleton was stained with 50 nM rhodamine phalloidin (Invitrogen) followed by the repeated wash in PBS. Then the coverslips containing treated cells were mounted with antifade reagent (Invitrogen) and allowed to dry for overnight. The fixed cells were observed under IX 81 microscope (Olympus).

### Modeling of F-actin ACD domain

The modelling of ACD domain containing WH2 motif with F-actin was carried out using the Cluspro server [45]. The blind docking was carried out using the three monomers taken from actin filament to mimic the F-actin as receptor and crystal structure of the VgrG-1-ACDof *Vibrio cholerae* (PDB Code:4DTL) as receptor [9]. We selected the final model on the basis of balanced score from the cluster with 37 members. Two clusters with WH2 domains participating in direct binding were shown in the images generated by Pymol [35]. The complex was further energy minimized by using AMBER 14 MD [46] simulation package to retain a low energy global conformation. The final model was then superimposed to the filaments to generate F-actin decorated ACD to show the binding of the VgrG-1-ACD to filamentous actin.

## Acknowledgement

PD thanks Indian Institute of Science Education and Research-Kolkata, India for providing her fellowship. This work was supported by Indian Institute of Science Education and Research-Kolkata.

## Author’s Contribution

Conceived and designed the experiments: SM SG AM. Performed the experiments: PD JAS MM DD. Analysed the data: SM SG AM. Wrote the paper: SM PD SG MM JAS and DD.

---

**Figure S1:**
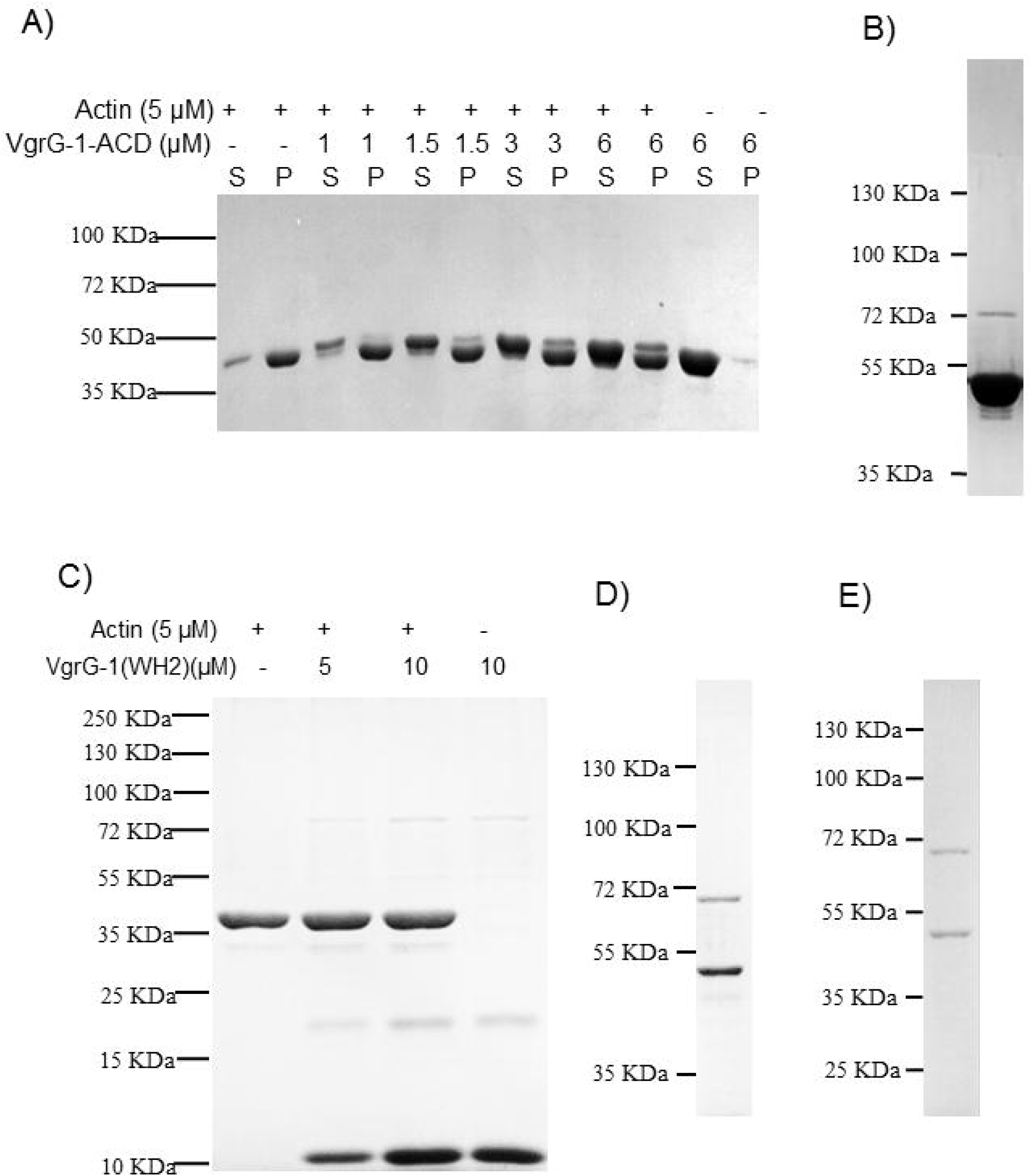
The VgrG-1-ACD binds to actin. **[A]** The co-sedimentation of VgrG-1-ACD with actin after the formation of actin filaments. **[B]** The I1017A& K1018A mutant ran in 10% SDS PAGE. **[C]** The wild type WH2 domain cannot cross link actin. **[D]** The V1013A & M1014A mutant purified as 6X His tag and ran in 12% SDS PAGE. **[E]** The V1013A & M1014A and I1017A& K1018A purified mutant protein ran in 12% SDS PAGE.

